# CLAPO syndrome: Identification of somatic activating *PIK3CA* mutations and delineation of the natural history and phenotype

**DOI:** 10.1101/154591

**Authors:** Lara Rodriguez-Laguna, Kristina Ibañez, Gema Gordo, Sixto Garcia-Minaur, Fernando Santos-Simarro, Noelia Agra, Elena Vallespín, Victoria E Fernández-Montaño, Rubén Martín-Arenas, Ángela del Pozo, Héctor Gonzalez-Pecellín, Rocío Mena, Inmaculada Rueda-Arenas, María V. Gomez, Cristina Villaverde, Ana Bustamante, Carmen Ayuso, Víctor L. Ruiz-Perez, Julián Nevado, Pablo Lapunzina, Juan C. Lopez-Gutierrez, Victor Martinez-Glez

**Affiliations:** Vascular Malformations Section, Institute of Medical and Molecular Genetics, INGEMM-IdiPAZ, Hospital Universitario La Paz, Madrid, 28046, Spain.; Bioinformatics Section, Institute of Medical and Molecular Genetics, INGEMM-IdiPAZ, Hospital Universitario La Paz, Madrid, 28046, Spain.; Clinical Genetics Section, Institute of Medical and Molecular Genetics, INGEMM-IdiPAZ, Hospital Universitario La Paz, Madrid, 28046, Spain.; Structural and Functional Genomics Section, Institute of Medical and Molecular Genetics, INGEMM-IdiPAZ, Hospital Universitario La Paz, Madrid, 28046, Spain.; Vascular Anomalies Center, Plastic Surgery, Hospital Universitario La Paz, Madrid, 28046, Spain.; CIBERER, Centro de Investigación Biomédica en Red de Enfermedades Raras, ISCIII, Madrid, 28029, Spain.; Instituto de Investigaciones Biomédicas “Alberto Sols”, CSIC-UAM, Madrid, 28029, Spain.; Department of Genetics, IIS-Fundación Jiménez Díaz UAM, Madrid, 28040, Spain.

## Abstract

**Background:** CLAPO syndrome is a rare vascular disorder characterized by Capillary malformation of the lower lip, Lymphatic malformation predominant on the face and neck, Asymmetry, and Partial/generalized Overgrowth. Although the genetic cause is not known, the tissue distribution of the clinical manifestations in CLAPO seems to follow a pattern of somatic mosaicism.

**Subjects and methods:** We clinically evaluated a cohort of 13 patients with CLAPO and screened 20 DNA blood/tissue samples from nine patients using high-throughput, deep sequencing.

**Results:** We identified five activating mutations in the *PIK3CA* gene in affected tissues from six of the nine patients studied; one of the variants (NM_006218.2:c.248T>C; p.Phe83Ser) has not been previously described in developmental disorders.

**Conclusions:** We describe for the first time the presence of somatic activating *PIK3CA* mutations in patients with CLAPO. We also report an update of the phenotype and natural history of the syndrome.

## Introduction

Vascular anomalies represent a broad spectrum of disorders produced by abnormal embryological development of blood vessels. The manifestations of these disorders may be prenatal, congenital, or postnatal. In general, they manifest with a gradual increase in size or extent, proportionally greater than the growth of the patient, resulting in a lesion that is more significant at puberty^1^. Vascular anomalies frequently occur as part of recognizable, pleiotropic developmental syndromes^2^, which includes overgrowth syndromes^3^.

CLAPO syndrome [OMIM: 613089] —Capillary Malformation of the lower lip, Lymphatic Malformation of the face and neck, Asymmetry and Partial/generalized Overgrowth — is a rare vascular disorder of unknown etiology first described in 2008 in six unrelated patients^4^. Only two additional patients have been described ^5^,^6^, although we suspect that the syndrome might be underdiagnosed. The tissue distribution of the clinical manifestations in patients with CLAPO is similar to other vascular and overgrowth syndromes caused by somatic mutations such as Proteus syndrome caused by activating mutations in *AKT1*^ref^ ^7^, Sturge Weber syndrome caused by activating mutations in *GNAQ*^8^, or PROS (*PIK3CA* Related Overgrowth Spectrum) caused by activating mutations in *PIK3CA*^9^.

Somatic mosaicism is defined as the presence of more than one clone of cells with different genotypes, all which are derived from a single cell. Mosaicism can contribute to variable clinical expressivity of a trait due to tissue-specific involvement. Consequently, the phenotypic spectrum of all these vascular and/or segmental overgrowth syndromes is heterogeneous and complex. This has become evident with the gradual inclusion of more syndromes and isolated malformations as mosaic, pathogenic variants are being detected in additional clinically distinct phenotypes, some of them sharing a common genetic cause.

Somatic mosaicism is a theoretical possibility in CLAPO, and even though it is a clinically well distinguished syndrome, it shares several clinical manifestations with previously mentioned syndromes as well as with other entities also caused by mutations in genes within the PI3K-Akt-mTOR pathway. Here we tested the hypothesis that CLAPO could be caused by somatic mutations activating this pathway, and we evaluated the natural history and phenotype of CLAPO and its phenotypic overlap with other PI3K-Akt-mTOR entities.

## Subjects and Methods

### Patients and samples

The study was performed at the Institute of Medical and Molecular Genetics (INGEMM) and the Vascular Anomalies Center, Plastic Surgery Department at La Paz Hospital, Madrid (Spain). We retrospectively reviewed the clinical characteristics and follow up of 13 patients diagnosed with CLAPO between February 2007 and March 2017, including the six patients reported in the original description of the syndrome ^4^. All patients were enrolled in the Spanish Overgrowth Syndrome Registry at La Paz Hospital. Clinical findings for all patients are summarized in Table I.

**Table I.** Phenotypic review of 13 patients with CLAPO.

| ID | Age | Sex | Years of follow-up | Biological samples | CM lower lip (HPO:0005306) | LM (HPO:0100766) | Overgrowth (HPO:0001548) | VM (HPO:0012721) | Other CM | Other |
| --- | --- | --- | --- | --- | --- | --- | --- | --- | --- | --- |
| P1 | 40 | Male | 10,2 | Blood and affected skin tissue | Full lip | Bilateral lymphedema with predominance of foot-knee | Partial overgrowth of the right side of the body (+1.5 cm.) | Multiple varicose veins in the legs | Right area of the body, reddish color | — |
| P2 | 17 | Female | 10,2 | Blood and affected skin tissue | 2 cm | Lower limbs | Left leg and right arm hypertrophy | — | Left inguinal region | — |
| P3 | 20 | Male | 9,4 | Blood | Full lip | Tongue and mandibular region | Generalized | Neck | — | — |
| P4 | 20 | Female | 10,1 | Blood and LM tissue | Full lip | Tongue and mandibular region | — | Neck | — | Pectus excavatum |
| P5 | 16 | Female | 10,0 | Blood | 4 cm | Tongue and mandibular region | — | Neck | — | — |
| P6 | 11 | Female | 10,2 | Blood and LM tissue | 2 cm | Tongue and mandibular region | — | Neck | Neck | — |
| P7 | 18 | Female | 2,6 | Blood, CM tissue and saliva | Full lip | Intraoral part of the lip | — | — | — | Multiple nevus |
| P8 | 12 | Male | 2,0 | Blood and affected mucosa tissue | 3 cm | Cervicofacial region, tongue, gluteus | — | Neck | Right gluteus | Left facial paralysis and hypotonia |
| P9 | 4 | Female | 1,5 | Blood and CM tissue | 3 cm | — | — | Neck | Multiple, extensive: right lower limb (genital and gluteal), face, neck | Presacral ganglioneuroma. |
| P10 | 2 | Female | 1,5 | Blood, CM tissue and saliva | Full lip | Cervicofacial region, tongue | Segmental hypoplasia | Neck | Left hand, between the 4th and 5th finger | Non-effective Rapamycin treatment. Ocular impairment due to LM. Gastrotomy. |
| P11 | 5 | Male | 1,1 | Blood and saliva | Full lip | Cervicofacial region | — | Neck | Glabella and neck | — |
| P12 | 4 | Male | 1,1 | Blood and saliva | Full lip | Cervicofacial region, tongue | — | Neck | Tongue and right ankle | — |
| P13 | 7 | Female | 0,8 | Blood and LM tissue | Gingiva, no lip involvement | Tongue and mandibular region | — | Tongue and gingiva. Possible VM in left hip | Hip | Propanolol not effective |

CM: Capillary malformation. LM: Lymphatic malformation. VM: Venous malformation. HPO: Human phenotype Ontology code.

Blood samples were collected from all patients, saliva samples from four patients and affected tissue samples from nine patients (Table I). Affected tissue was defined as any type of vascular malformation obtained during surgical procedure scheduled as part of the routine treatment, and in the case of cutaneous biopsies, these were obtained in regions with vascular malformation and/or true overgrowth (differentiating it from an asymmetry as a consequence of a vascular malformation). DNA extraction was performed by standard procedures. All studies in this project were approved by the Ethics Committee of the Hospital Universitario La Paz (PI-1919), with informed consent.

### High-throughput sequencing

High-throughput sequencing studies were performed using three distinct custom panels previously available and validated for use in our laboratory. The subsequent analyses of the sequencing data were based on the use of virtual panels (i.e., tiers) with an average expected reading depth of 500X. The first panel (Tier1: Mosaic panel) included 20 genes known to cause developmental syndromes in the form of somatic mosaicism. The second panel (Tier2: OGLYVAS-OverGrowth, Lymphatic and Vascular Syndromes panel) was composed of 301 genes associated with vascular malformations and overgrowths of vascular origin, including PI3K-Akt-mTOR pathway genes, as well as 20 genes of the Mosaic panel. The third panel (Tier3: EMG) included 1,375 genes, including genes in OGLYVAS panel. The complete lists of genes can be found in the **Supplementary Table I**.

The three panels were designed, captured, and analyzed with the same tools, protocols, and instruments. The custom panels were designed by using NimbleDesign software (Roche NimbleGen, Inc. USA): hg19 NCBI Build 37.1/GRCh37.p13, targeting >98% of all exons (RefSeq) for these genes. For each sample, paired-end (2x150 bp reads) libraries were created according to the standard deep sequencing protocols KAPA HTP Library Preparation Kit for Illumina platforms, SeqCap EZ Library SR (Roche NimbleGen, Inc. USA) and NEXTflex-96 Pre Capture Combo Kit (Bioo Scientific) for indexing. The captured DNA samples were sequenced on a NextSeq™ 500 instrument (Illumina, Inc. USA) using a HIGH v2 300 cycles cartridge, according to the standard operating protocol.

### Bioinformatics analysis pipeline

Data generated by the NextSeq™ 500 Desktop Sequencer was analyzed using an in-house bioinformatics pipeline for somatic mosaicism detection. BCL files containing base calls were converted into paired FASTQ files using bcl2fastq-v2.15.0.4 software from Illumina (https://github.com/brwnj/bcl2fastq) and pre-processing using Trimmomatic^10^ for trimming and cropping FASTQ data as well as removing adapters. Subsequently, balanced reads were mapped to hg19/GRCh37 human genome by using Bowtie2 aligner^11^, and PCR duplicate reads were removed using Picard MarkDuplicates (http://broadinstitute.github.io/picard/). A subsequent local realignment and recalibration of reads was done to correct misalignments due to the presence of INDELs by using GATK RealignerTargetCreator and IndelRealigner, and BaseRecalibrator methods respectively. Once this pre-processing was achieved, the flowchart depicted in **Supplementary Figure 1** was followed.

The initial specific approach for mosaic detection included the extraction of the base-pair information for each genomic position from the BAM files using samtools mpileup v1.3^ref^ ^12^, facilitating the subsequent SNP/INDEL calling. The base quality (q value) and the mapping quality scores (Q value) were lowered to 0, to characterize mutated alleles amplified in very low proportion and to avoid losing variants in non-unique genomic sequences, since many genes in the capture design have pseudo-genes or share high percent identity with other genomic sites.

Variant calling was performed using bcftools v1.3. The low constraints allowed the emergence of many alternate alleles, even though many could be false positives. The strategy of the analysis was to keep all multiallelic sites in the VCF file for later consideration. Some attributes were defined to filter out likely sequencing artifacts or variants with high frequency in samples sequenced in the same run, and others to keep mosaic variants to analyze further on. This was applied in all the samples individually, regardless of the tissue type, generating a corresponding VCF file (see **Supplementary Figure 1a**).

Subsequently, a germ line versus somatic variant comparison was undertaken by analyzing the tissue and the blood VCF files from the same patient/sample (see **Supplementary Figure 1b**). The contrasting of matched tissue and blood samples was crucial to distinguish somatic from germline variants, bringing out low allelic fraction from background noise caused by the high-throughput sequencing error rate. The final files encoded global information about alignments. Manual filtering was applied to determine the candidate pathogenic variants. The resulting VCF files were manually visualized with the Integrative Genomics Viewer (IGV) program to verify mutations and correct annotation.

### Validation of high-throughput deep sequencing variants (Sanger, pyrosequencing, and ddPCR)

**Sanger sequencing** was used to confirm non-mosaic heterozygous variants and variants present in more than 15% of the reads in the deep sequencing data. Standard PCR and Sanger sequencing were performed using the 96-capillary ABI 3730xl ADN analyzer (Applied Biosystem, Foster, USA). Long range PCR and Sanger sequencing were also used to verify chromosomal location in variants found with high identity to another part of the genome. **Pyrosequencing** primers were designed using PyroMark software, and QIAGEN reagents and the Pyromark Q96 MD instrument (QIAGEN, USA) were used according to manufacturer’s protocol to confirm mosaics variants in the 5% and 15% read fraction range. **Droplet digital polymerase chain reaction** (ddPCR), able to detect and quantify somatic mosaic variants at frequencies as low as 0.1%^13^, was used to detect/validate and quantify variants found on deep sequencing studies in less than 5% of reads or variants above 5% of reads but whose confirmation by pyrosequencing was not conclusive. We also used ddPCR to screen for the three common mutations described in PROS (*PIK3CA*: NM_006218.2: c.1624G>A, p.Glu542Lys; c.1633G>A, p. Glu542Lys; and c.3140A>G, His1047Arg) in all samples in which no variants were previously detected by high-throughput deep sequencing ^14^. The QX200 Droplet Digital PCR System (Bio-Rad) was used to generate 20,000 DNA-oil droplets according to the manufacturer’s protocol. Nucleic acid molecules were quantified using a two-color fluorescence detector (FAM channel for mutant allele and VIC channel for wild type) and six assays with well-defined and validated somatic mosaic variants were used as controls to validate the technique.

## Results

### Clinical Features

Longitudinal data of 13 patients with CLAPO was gathered for a total of 70.5 person-years (ranging from 0.8 to 10.2). The mean age at diagnosis was 8.0 years (0.6 to 30.4) and the mean age at the end of this study was 10.4 years (2.2 to 40.5). None of the patients had a relevant family history. Except for one patient of Arab descent (P6), all patients in the cohort were Caucasian and had parents from the Iberian Peninsula. The CLAPO cohort included eight females and five males. Among the main features of CLAPO the only common clinical manifestation was the capillary malformation (CM) of the lower lip (13/13; 100%); all other features were present at birth or became evident postnatally. The CM of the lower lip was always present in the midline, with a symmetrical distribution, ranging from 2 to 11 cm, with well-defined borders in several patients, and often affecting the portion of the skin under the lip or the intra-oral mucosa. We observed three distinct patterns of midline CM on the lower lip: 1) five patients (P2, P5, P6, P8, P9) had a narrow midline CM with reddish color, 2) seven patients (P1, P3, P4, P7, P10, P11, P12) had a CM that involved the entire lower lip with a predominant brown/purple color and 3) one patient (P13) had a CM including the central inner area of the lower lip without external skin involvement under the vermilion. In our experience, untreated isolated capillary malformations (classically named port-wine stains) evolve by a progressive thickening and darkening of the malformation due to a progressive vessel dilation, which can, in turn, produce skin overgrowth. However, after ten years of follow-up we did not observe this type of progression, but we did observe decreased intensity in the CM of the lower lip in four of six patients followed more than nine years. While mid-line CMs of the face are frequently correlated with lesions of the brain in patients with Megalencephaly-Capillary malformation syndrome^15^, no psychomotor delay or intellectual deficit was apparent in CLAPO patients.

The second major clinical feature was the presence of lymphatic malformations (LM), observed in 12 of 13 patients (92%). In 10 patients (P3-8, P10-13) the LM involved the lip, oral mucosa, neck, and tongue, seen in five of ten with right sided predominance (P5, P6, P10-12). Other two patients (P1, P2) had unilateral LM on the lower limbs, one of which (P1) was associated with lymphedema. At birth, the tongue had a symmetrical midline combined capillary/lymphatic/venous malformation in eight of 13 patients (P3-6, P8, P10, P12, P13) with CLAPO, which was mild, well-defined, and affected the tip of the tongue. In five of seven patients (P3-5, P8, P13), the tongue lesions evolved over time, causing growth of the affected area and severe hemorrhagic events with episodes of acute inflammation, triggering and aggravating symptoms. On the other hand, Patient 2 had a left thigh LM that was not noted until 15 years of age, indicating that an apparent absence of congenital LM does not exclude a later onset. Sporadic LMs are rare and arise more often with craniofacial venous malformations (VM)^16^. In our cohort LMs of the neck, tongue, and/or limb were also observed in association with VM in eleven patients (all but patients 2 and 7), which showed progressive increase of size and in no case involution.

In this cohort the asymmetry in eight of 13 patients (P4-6, P8, P10-13) actually reflected a direct consequence of the presence of a LM on face/neck, although in three patients (P1-3) there was a true asymmetric overgrowth with the presence of bony hypertrophy. There was also one patient (P10) in which asymmetry was caused by left limb undergrowth. Patient 1 had macrodactyly and overgrowth of the right side of the body, and Patient 2 had a postnatal increase in the length of the left leg.

Recently, patient 9 suffered a large presacral ganglioneuroma that was successfully resected, and patient 10 had severe failure to thrive needing gastrostomy. Those findings can eventually expand the phenotype of CLAPO syndrome, as we collect more clinical information and investigate a larger cohort of patients with CLAPO.

### Molecular results

Our main hypothesis was that somatic mutations were the genetic cause of CLAPO. Accordingly, we screened 20 DNA blood/tissue samples from nine patients with CLAPO using custom high-throughput deep sequencing panels including genes associated with vascular malformations and overgrowth, and a custom bioinformatic pipeline for somatic mosaic detection with a tiered analysis approach. Average read depth for Tier1 was 535X, for Tier2 342X, and for Tier3 404X, after removing duplicates. Variants were excluded as disease candidates by their presence in >0.01 population frequency (1000 Genomes project (http://www.internationalgenome.org/), NHLBI Exome Variant Server (http://evs.gs.washington.edu/EVS), Exome Aggregation Consortium (ExAC; http://exac.broadinstitute.org)), by pathogenicity predictors, and by using new custom parameters developed here for somatic mosaic detection (AVAF, SampleRun). After filtering, mean variants per sample for Tier1 was 5.2, for Tier2 31.3, and for Tier3 175.6.

Our analysis identified *PIK3CA* variants in six of nine (75%) patients with CLAPO (P1, P2, P6, P9, P10, and P13). All mutations were somatic missense single nucleotide variations (SNV) with a range of mosaicism between 1.26% and 16%. We identified a total of five distinct mutations in *PIK3CA* (NM_006218.2), including one hotspot mutation (c.1624G>A;p.Glu542Lys), two recurrent, strong (gain of function) mutations (c.3140A>T;p.His1047Leu, c.1258T>C;p.Cys420Arg), one previously described mutation in patients with macrodactyly (c.344G>C;p.Arg115Pro), and one novel somatic *PIK3CA* mutation (c.248T>C; p.Phe83Ser) not previously described in developmental disorders.

All variants detected in tissue samples were confirmed using at least one orthogonal method based on the mosaicism percentage of the alternative allele from the NGS candidate gene sequencing. None of the mutations detected in the patients were present (by Sanger sequencing) in the blood samples of the patient’s parents. Molecular data and variant information were collected in Figure 1.

**Figure 1.**
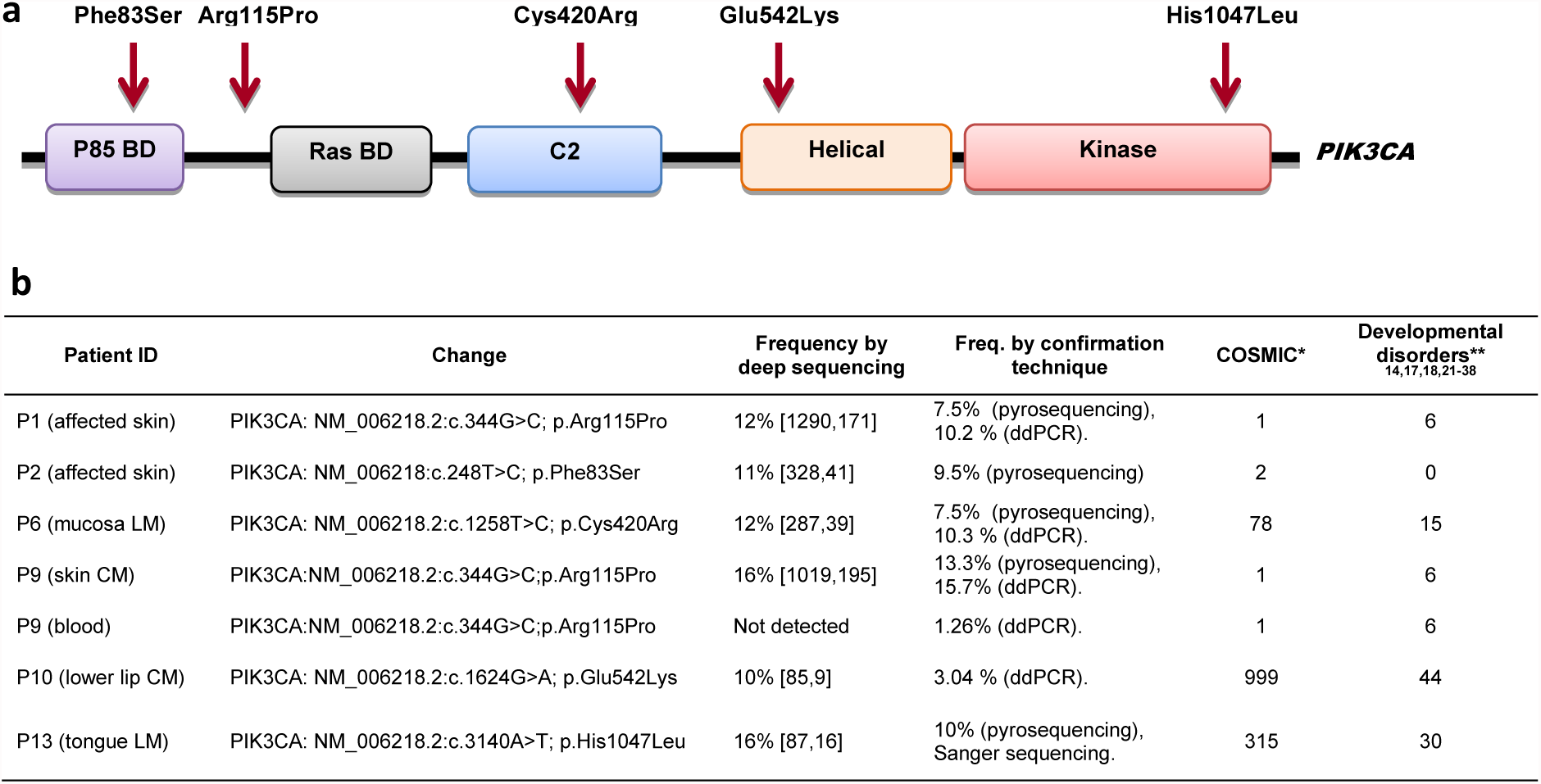
*PIK3CA* mutations detected in CLAPO patients. **a:** Diagram of the distribution of the mutations detected along the *PIK3CA* domains gene. **b:** Information about each tissue specific variant detected in nine patients with CLAPO. Variant frequency by deep sequencing shows the percentage of mosaicism and the number of reads for each allele (wt, alternate). Frequency by confirmation technique includes which other technique/s were used to validate the candidate variant and its percentage. * Cancer samples with identified *PIK3CA* mutations in COSMIC database (http://cancer.sanger.ac.uk/; accessed June 2017). ** Number of patients with vascular/overgrowth disorders previously reported with the specific *PIK3CA* variant.

## Discussion

In the present report we describe for the first time the presence of somatic activating *PIK3CA* mutations in patients with CLAPO, and we report an update of the phenotype and natural history of the syndrome.

Somatic mosaic mutations in *PIK3CA* have been detected in six of nine patients with CLAPO syndrome. All mutations were detected in affected tissue, in percentages ranging from 3 to 16%, and in one of the patients (P9) the mutation was also mosaic at a level of 1.26% in a blood sample. All the mutations detected are missense variants, four of which have previously been reported as causing the PROS spectrum^9^,^14^ and detected in different types of cancer (COSMIC database. http://cancer.sanger.ac.uk/). We also identified a novel *PIK3CA* mutation (Phe83Ser), not previously described in development disorders, but then again reported in the COSMIC database in two cancer samples.

The family of syndromes caused by activating *PIK3CA* mutations includes well-known and distinct entities such as Megalencephaly-Capillary malformation (MCAP), CLOVES syndrome (Congenital Lipomatous Overgrowth, Vascular malformations, Epidermal nevi, And Skeletal/Spinal abnormalities), isolated lymphatic malformations (LM), isolated adipose fibrodysplasia and hemimegalencephaly, among others^9^. In addition to the distribution pattern, some of the syndromes included in PROS have overlapping clinical findings with CLAPO, mainly capillary and lymphatic malformations and segmental overgrowth. However, sharing a common mutated gene does not make their formerly differentiated classification obsolete since it has implications for clinical diagnosis, evolution, and follow-up.

Our findings suggest that somatic activating *PIK3CA* mutations can be responsible for the combination of capillary, lymphatic, and venous malformation as well as the overgrowth features characteristic of CLAPO syndrome. Given the overlapping phenotypic features between CLAPO and PROS (mainly with MCAP and CLOVES), and the presence of somatic activating *PIK3CA* mutations, we postulate that CLAPO syndrome might be included into the PROS spectrum.

The multiple or reticulate and diffuse capillary malformations present in different parts of the body appear to be similar in both CLAPO and MCAP. However, one of the main clinical manifestations in CLAPO is the capillary malformation of the lower lip, a feature that is not considered characteristic of MCAP, in which midline CMs are most frequently described in upper lip, glabella, and philtrum. Anatomically, the differential involvement of the lower lip or the upper lip is distinctive and is relevant as they have distinct embryological origins and clinical implications. The forehead, the dorsum apex of the nose, and the central part of the upper lip derive from the frontonasal prominence, whereas the chin, lower lip, and lower cheek regions derive from the mandibular prominences. These distinct embryological origins might also explain the co-occurrence of brain alterations and CM of the upper lip in MCAP and the absence of neurological involvement in CLAPO.

The second main feature in CLAPO, the lymphatic malformation, is also frequent in CLOVES and facial infiltrating lipomatosis. While two of the 13 patients described here had a lymphatic malformation of the legs, this type of malformation occurs in CLAPO more frequently in the cervicofacial region and tongue, where there is a clear non-random association with capillary or venous malformations. Venous malformations were not originally described as a main feature in CLAPO, but we now know that it is a rather frequent feature, and it has recently been described that this type of malformations can arise sporadically as a consequence of somatic activating mutations in *PIK3CA*^17^, which has also been described for isolated lymphatic malformations^18^. Since it is possible that both lymphatic and venous malformations are not apparent congenitally, the presence of capillary malformation in the lower lip should make us aware of a possible later appearance of these two types of vascular malformations. This specific and non-random combination of vascular malformations highlights the utility of the clinical diagnosis in patients with CLAPO.

Overgrowth is not always evident in CLAPO, as none of the patients had macrocephaly, the involvement was segmental and rarely generalized, and in the case of facial asymmetry it appears to be caused by a vascular component. As with all overgrowth syndromes, it is important to distinguish in CLAPO and PROS what is apparent asymmetry from true overgrowth caused by hyperplasia or hypertrophy. In spite of the fact that partial/generalized overgrowth is part of the CLAPO acronym, its low frequency in this cohort of patients suggests that it should not be considered a major clinical criteria for the diagnosis of CLAPO, despite being a frequent (but not mandatory) feature in PROS.

Considering the clinical characteristics and the natural history described in our cohort of patients, CLAPO could be classified as an entity with intermediate severity in the PROS spectrum, sharing characteristics mainly with MCAP and CLOVES, and certainly with isolated forms of lymphatic and venous malformations.

One of the major challenges in the growing number of syndromes caused by alterations in the form of somatic mosaicism is the detection of altered alleles in low mosaics in affected tissues. Correspondingly, the molecular diagnosis of CLAPO patients is not trivial. One limitation comes from the requirement to obtain affected tissue in order to carry out the detection of the pathogenic variants. Thus, the lack of success detecting *PIK3CA* mutations in three of nine of the molecularly studied patients with CLAPO does not implicate the absent of mutations in this gene, neither excludes the molecular diagnosis of CLAPO.

Another challenge is the bioinformatic method for the detection of somatic mosaicism. There are several computational approaches to detect somatic variants, but they cannot be applied to this problem since the starting data or the genomic landscape is different. Strelka^19^ for SNV discovery or Mutect^20^ for INDEL discovery are two of the most used variant callers for somatic variant detection. However, neither of them was developed to detect variants in mosaic disorders, which in the case of *PIK3CA* is also complicated by the fact that exons 10 to 14 (chromosome 3) have a 98% identity with another region of the genome located in chromosome 22. Thanks to the computational approach applied here, we could rescue reads with low mapping quality score due to high percent identity with other genomic regions, and call variants that were present both in somatic and mosaic state.

In conclusion, this study documents the clinical features and natural history in a well-defined cohort of patients with CLAPO, and detected activating *PIK3CA* mutations in six of nine studied affected tissues. We detected *PIK3CA* activating variants previously described as causing the PROS spectrum, together with a clinical distribution pattern distinctive of somatic mosaicism, and a constellation of non-random clinical manifestations that, although in combination are specific to CLAPO syndrome, are also frequent in other syndromes within the PROS spectrum. We conclude that CLAPO belongs to the PROS family of somatic syndromes. In clinical terms, the differences between CLAPO and other PROS spectrum phenotypes demonstrates that the presence of a CM in the upper lip should raise suspicion of possible neurological involvement, whereas the CM in lower lip is associated with later occurrence of associated lymphatic/venous malformations. In diagnostic terms, the approach in CLAPO requires the use of appropriate samples and bioinformatic algorithms, allowing the detection of somatic mosaic variants. Because CLAPO is caused by activating mutations in *PIK3CA*, affected patients may benefit from inhibitors of this pathway. Therapeutic trials of this disorder should be undertaken.

## Acknowledgements

We are grateful to the patients and their families for their participation in this study. We want to honor the memory of Angel LLopis and thank his father Fernando. This work has been partly possible thanks to them. We also would like to thank Dr. Leslie Biesecker of the National Human Genome Research Institute, NIH, USA, for his invaluable contribution revising this work critically.

## Declarations

Ethics approval and consent to participate: Written informed consent was obtained from all patients/families to participate in this study, and the project was approved by the ethical committee of the Hospital Universitario La Paz (reference PI-1919). Consent to publish: Consent for publication was obtained from all patients and/or parents. Availability of data and material: The datasets used and/or analyzed during the current study are available from the corresponding author on reasonable request. Competing interests: All authors declare no competing interests. Funding: This research was supported by the project “Genetics of vascular and lymphatic malformations” financed with funds donated by Asociación Ultrafondo and Villareal FC, and Co-financed by the project IP-17 from the call “Todos Somos Raros” (Telemaraton TVE promoted by Fundación Isabel Gemio, Federación ASEM, and Federación Española de Enfermedades Raras).

